# Sex determination through X-Y heterogamety in *Salix nigra*

**DOI:** 10.1101/2020.03.23.000919

**Authors:** Brian J. Sanderson, Guanqiao Feng, Nan Hu, Craig H. Carlson, Lawrence B. Smart, Ken Keefover-Ring, Tongming Yin, Tao Ma, Jianquan Liu, Stephen P. DiFazio, Matthew S. Olson

**Author notes:** Department of Biology, West Virginia University, Morgantown, WV, 26506-6057 USA.

## Abstract

The development of non-recombining sex chromosomes has radical effects on the evolution of discrete sexes and sexual dimorphism. Although dioecy is rare in plants, sex chromosomes have evolved repeatedly throughout the diversification of angiosperms, and many of these sex chromosomes are relatively young compared to those found in vertebrates. In this study, we designed and used a sequence capture array to identify a novel sex-linked region (SLR) in *Salix nigra*, a basal species in the willow clade, and demonstrated that this species has XY heterogamety. We did not detect any genetic overlap with the previously characterized ZW SLRs in willows, which map to a different chromosome. The *S. nigra* SLR is characterized by strong recombination suppression across a 2 MB region and an excess of low frequency alleles, resulting in a low Tajima’s D compared to the remainder of the genome. We speculate that either a recent bottleneck in population size or factors related to positive or background selection generated this differential pattern of Tajima’s D on the X and autosomes. This discovery provides insights into factors that may influence the evolution of sex chromosomes in plants and contributes to a large number of recent observations that underscore their dynamic nature.

## Introduction

Some groups of animals exhibit surprisingly frequent movement in the genomic locations of sex determination loci (Gammerdinger and Kocher, 2018; Miura, 2007), but it is not yet clear whether this is common in plant species, where the evolution of separate sexes (dioecy) is more frequent compared with most animal groups (Beukeboom and Perrin, 2014). In mammals (Cortez *et al*, 2014; Lahn and Page, 1999) and birds (Ellegren, 2010; Shetty *et al*, 1999; but see Sigeman *et al*, 2019), most species have maintained similar sex chromosomes for over 80-130 My. In other animal groups, however, sex chromosome transitions appear more common. For instance, among several genera of cichlid fishes the sex determination regions reside on 12 different sex chromosomes, with both male and female heterogametic systems (XY and ZW, respectively; Gammerdinger and Kocher, 2018). Similarly, Dipteran flies harbor more than six different sex chromosomes, and multiple sex chromosome transitions have occurred within the genus *Drosophila* (Vicoso and Bachtrog, 2015). However, in plants few taxonomic groups that share the same origin of dioecy have been studied in detail (Charlesworth, 2002; Charlesworth, 2016a). Two notable exceptions are octoploid *Fragaria* (strawberry), where a cassette of sex determining genes has moved among homeologs, acquiring linked genomic regions as it moved, likely all within the last 1 My (Tennessen *et al*, 2018) and *Populus*, where sex determination genes have moved and switched from male to female heterogamety resulting from changes in the regulation of an *ARR17* ortholog, a type-A cytokinin response regulator (Muller *et al*, 2020; Yang *et al*, 2020). In this regard, the willows (genus *Salix*) are of particular interest because they are exclusively dioecious and diverged from a their sister genus *Populus* in the mid-Eocene, 40-50 million years ago (Berlin *et al*, 2010; Zhang *et al*, 2018).

Chromosome 15 was originally identified as the location of the sex-linked region (SLR) in *Salix* using sex-linked SCAR markers (Alstrom-Rapaport *et al*, 1998; Temmel *et al*, 2007), which was later confirmed and characterized as ZW heterogamety in three species of willow from subgenus Vetrix using genome resequencing and mapping (*Salix viminalis*: (Pucholt *et al*, 2015); *S. suchowensis*: (Hou *et al*, 2015); and *S. purpurea*: (Zhou *et al*, 2018)). The most intensively studied of these species is *S. purpurea*, in which the SLR is a 6.7 MB nonrecombining region (Zhou *et al*, 2020; Zhou *et al*, 2018). The location and pattern of heterogamety of the SLR in these species of *Salix* differ from those found in the sister genus *Populus*, where the SLR commonly exhibits XY heterogamety and is located on chromosome 19 (Geraldes *et al*, 2015; Pakull *et al*, 2011; Pakull *et al*, 2009).

Genetic diversity on sex chromosomes and autosomes is shaped by both demographic and selective factors (Charlesworth *et al*, 1987; Ellegren, 2009; Vicoso and Charlesworth, 2006). The expected effective population size (N_e_) of the X chromosomes is ¾ that of the autosomes, resulting the stronger impacts of genetic drift on the X than on autosomes and a general expectation that diversity on the X should be ¾ that of diversity on autosomes (Ellegren, 2009; Sayres, 2018). At demographic equilibrium, however, the site frequency spectrum (SFS) for neutral alleles should be similar for both the X chromosomes and autosomes, resulting in similarity for statistics such as Tajima’s D which is sensitive to the frequency of alleles in the population (Tajima, 1989). However, because of its lower N_e_, the X is more strongly impacted by bottlenecks in population size resulting in greater reductions in Tajima’s D for the X than for autosomes (Gattepaille *et al*, 2013). Also, because X chromosomes spend less time in males than in females, differences in the variance of reproductive output and generation time between the sexes can differentially influence diversity on the X and autosomes (Amster and Sella, 2020; Charlesworth, 2001). Because the parts of the X chromosome aside from the pseudo-autosomal regions (PARs) do not recombine in males, LD is often elevated compared to autosomes. Thus, sites on the non-recombining regions of the X chromosome are often more strongly affected by hitchhiking with sites that are directly impacted by either positive or purifying selection (Betancourt *et al*, 2004; Charlesworth *et al*, 1987). Also, although recessive or partially recessive alleles on autosomes can be sheltered from the effects of selection in females, because the X chromosome is hemizygous in males, alleles are exposed to selection (Charlesworth *et al*, 1987; Haldane, 1924) resulting in a stronger response to selection for X-linked genes than is typical for autosomal genes (Vicoso and Charlesworth, 2006).

The goals of the present study were to investigate the location and heterogamety of the SLR in *Salix nigra*, a species in a subgenus of *Salix* within which sex chromosomes have not yet been mapped, and to investigate patterns of diversity in its SLR. This species is a tree-form willow in the subgenus Protitea that is basal in the *Salix* lineage (Argus, 1997; Barkalov and Kozyrenko, 2014). Using targeted sequence capture to selectively genotype 24 males and 24 females with genome-wide markers, we show that a region on chromosome 7 exhibits highly divergent genotypes between males and females, which indicates that this interval harbors the SLR. A comparison of heterozygosity between males and females, as well as the identification of a pattern of slow decay of linkage disequilibrium among males in this region is consistent with an XY sex determination system, which we confirmed with amplicon sequencing. This is the first evidence that the location and heterogamety of the sex chromosomes are variable in the genus *Salix*. We also found a greater proportion of low frequency alleles in the SLR compared to the autosomes, which we argue likely results from either a recent selective sweep or rare recombination with the Y chromosome combined with background selection on the X.

## Methods

### Probe Design

To consistently genotype common sites in a reduced representation of the genome, we designed a targeted sequence capture array (RNA probe hybridization) to efficiently capture exonic regions throughout the genome across species in the genus *Salix*. As target regions diverge due to sequence polymorphism both the efficiency of RNA probe binding and capture efficiency are reduced (Lemmon and Lemmon, 2013). Thus, we quantified sequence polymorphism among whole-genome resequencing data from a diverse array of *Salix* species, including *S. arbutifolia, S. bebbiana, S. chaenomeloides, S. eriocephala, S. interior, S. scouleriana*, and *S. sitchensis*. Whole-genome short reads of the *Salix* species were aligned to the *S. purpurea* 94006 genome assembly version 1 (Salix purpurea v1.0; Carlson *et al*, 2017; DOE-JGI, 2016; Zhou *et al*, 2018) using bwa mem (Li, 2013), single nucleotide polymorphisms (SNPs) and insertion-deletion mutations (indels) were identified using samtools mpileup (Li, 2011), and read depth for the variant calls was quantified using vcftools v. 0.1.15 (Danecek *et al*, 2011). A custom Python script was used to identify variant and indel frequencies for the aligned species at all exons in the *S. purpurea* genome annotation.

We further screened candidate regions to exclude high-similarity duplicated regions by accepting only loci with single BLAST hits against the highly contiguous assembly of *S. purpurea* 94006 version 5 (Zhou *et al*, 2020), which is less fragmented than the *S. purpurea* 94006 version 1. For the remaining loci, we selected regions across which at least 360 base pairs (bp) of exon sequence contained 2-12% difference in single nucleotides and fewer than 2 indels across all aligned species. For a majority of these genes, we identified multiple regions within each exon. These candidate regions were sent to Arbor Biosciences (Ann Arbor, MI, USA) for probe synthesis. Probes were designed with 50% overlap across the targeted regions, so that each targeted nucleotide position would potentially be captured by two probes. The final capture array consisted of 60 000 probes that target exonic sequences for 16 580 genes (Table S1). This array is available from Arbor Biosciences (Ref#170623-30).

### Library Preparation and Sequence Capture

The sex of 24 males and 24 females of *S. nigra* was identified from flowering catkins in a wild population near Dickens Springs, TX, USA in April 2017. Leaf tissue was collected and dried on silica beads. DNA was extracted from leaf tissue using the Qiagen DNeasy Plant Minikit (Qiagen, Hilden, Germany) and fragmented using sonication with the Covaris E220 Focused Ultrasonicator (Covaris, Inc., Woburn, MA, USA). Libraries were prepared using the NEBNext Ultra II DNA Prep Kit (New England Biolabs, Ipswitch, MA, USA), and quantified using an Agilent Bioanalyzer 2100 DNA 1000 kit (Agilent Technologies, Santa Clara, CA, USA). Libraries were pooled at equimolar concentrations into sixteen pools of six prior to probe hybridization to targeted capture probes following the Arbor Biosciences myProbes protocol v 3.0.1. The hybridized samples were subsequently pooled at equimolar ratios and sequenced at the Oklahoma Medical Research Foundation (Oklahoma, OK, USA) using a HiSeq 3000 (Illumina, Inc., San Diego, CA, USA).

### Variant calls and hard filtering

Reads were aligned to the version 5 assembly of *Salix purpurea* 94006 (female, ZW), using bwa mem (Li, 2013). We estimated the depth of read coverage across all targeted genes using bedtools intersect v. 2.25.0 (Quinlan and Hall, 2010). The alignments were screened for optical duplicates following the Broad Institute’s best practices recommendations (DePristo *et al*, 2011). Variants were identified using HaplotypeCaller in GATK v4.0.8.1 (McKenna *et al*, 2010) and initially called separately for each individual as GVCFs. The GVCFs were merged and variant sites and indels were called for all individuals using GenotypeGVCFs in GATK. Based on the distribution of quality scores, variant sites were screened with the hard filters: MQ < 40.00 ‖ SOR > 3.000 ‖ QD < 2.000 ‖ FS > 60.000 ‖ MQRankSum < -12.500 ‖ ReadPosRankSum < -8.000 ‖ ReadPosRankSum > 8.000. Individual genotypes were filtered out if supported by < 6 reads, or > 125 reads. Finally, genotypes that did not pass filtering criteria were assigned as no call genotypes (./.).

### Location of the sex determination region

To determine the genomic regions exhibiting the greatest differences between males and females, we performed a genome-wide association study (GWAS) in which we regressed sex (male or female) on the filtered variants using the program emmax (Kang *et al*, 2010) with a Bonferroni-adjusted significance threshold of *P* < 0.05 to assess the significance of sex linkage. The results were visualized with a Manhattan plot using the R package ggplot2 (Wickham, 2009). We quantified heterozygosity of genotypes at the loci identified with significant sex linkage directly from the filtered VCF file using the program vcftools v. 0.1.15 (Danecek *et al*, 2011).

To confirm the sex association, we analyzed an independent population of 16 male and 16 female *S. nigra* trees collected from the vicinity of Morgantown, WV. Primers were designed in conserved regions that flanked two loci that showed significant sex-linkage in the Texas population (Table S2). These were sequenced using Sanger chemistry on an ABI3130XL sequencer. Base calls were made using PHRED (Ewing and Green, 1998) and sequences were assembled using PHRAP and Consed (Gordon *et al*, 1998). Polymorphisms were identified using PolyPhred (Nickerson *et al*, 1997) and confirmed by visualization in Consed. The frequency of heterozygosity for males and females was compared for all loci with minor allele frequency > 0.35, which effectively screened out mutations that may have arisen since establishment of the SLR, as well as any instances of genotyping or sequencing error.

### Population genomics statistics

Kinship was estimated using relatedness2 in vcftools, which applies the methods used in KING (Manichaikul *et al*, 2010). Linkage disequilibrium (LD) for all pairs of loci was calculated as the squared allele frequency correlation (*r*^2^) using vcftools v. 0.1.15 (Danecek *et al*., 2011). LD decay was calculated across the SLR in males and females as well as the upstream pseudo autosomal region PAR1 from the end of chromosome 7 to 1 MB from the beginning of the SLR (PAR1: 0.1 MB – 3.88 MB) and from 1MB downstream of the SLR to the end of PAR2 (7.88MB - 13.0 MB) using formulas in Hill and Weir (Hill and Weir, 1988) and Remington *et al*. (Remington *et al*, 2001). The 1MB gaps at the beginning and end of the SLR were to avoid regions close to the edges of the SLR than may have intermediate recombination rates. To compare genetic diversity within the *S. nigra* SLR to diversity in other regions of the genome, we calculated per-site nucleotide diversity as π(Nei and Li, 1979) using vcftools v. 0.1.15 (Danecek *et al*, 2011) after filtering for only biallelic SNPs. We calculated πseparately for males, which carry both X and Y chromosomes, and females, which carry two X chromosomes. The average πfor 5 Kb sliding windows was calculated using R (R Core Team, 2018). To assess whether non-neutral processes may have influenced the SLR, we calculated values of Tajima’s D (Tajima, 1989) using the sliding window method implemented in vcftools v. 0.1.15 (Carlson *et al*, 2005; Danecek *et al*, 2011), with a window size of 25 Kb. Windows with fewer than 5 segregating sites were removed from the Tajima’s D analyses.

The scripts described above as well as the full details of these analyses including the alignment and variant calling pipeline are available in Jupyter notebooks at https://github.com/BrianSanderson/salix-nigra-slr.

## Results

The sequence capture reads aligned to an average of 47.82 ±1.47 (mean ± sd) Mb of the *S. purpurea* reference genome among the 48 libraries before filtering (approximately 14.5% of the 329.3 Mb genome). This corresponded to an average read depth of 6.17 ± 2.06X (mean ± sd) at sites genome-wide, and an average read depth of 44.68 ± 2.68X (mean ± sd) at on-target sites across the 16 580 targeted genes (Tables S1 & S3). The sequence capture probe set resulted in relatively even coverage across the genome (Fig S1, although unsequenced blocks of several thousand kb were present throughout the genome. This was especially true for the region close to the likely centromere on chromosome 7. Some regions also were sequenced that were not targeted (Fig S1). These off-target captures likely resulted from regions immediately up- and downstream of probe targets, as well as regions with weak homology to probes. A total of 6 826 811 SNPs was identified among the 48 individuals, of which 2 011 511 passed the filtering criteria, for an average SNP density in the sample of 95 SNPs/kb. The average kinship among individuals in the population was 0.176, indicating an average degree of kinship between half-sibs and first cousins in this population (Fig. S2) and suggesting that overall diversity estimated in this population may not be representative of the species as a whole.

A total of 38 SNPs on chromosome 7 between 4.88 Mb and 6.81 Mb exhibited significant associations with sex (P_*FDR*_ < 0.05; Figs. 1 & S3; Table S4). We refer to this region as the sex-linked region (SLR). An additional 2 SNPs on scaffold 197 and 5 SNPs on scaffold 257 also exhibited significant associations with sex (P_*FDR*_ < 0.05; Figs. 1 & S3; Table S4). We suspect that with additional data these scaffolds will be assembled to the SLR on chromosome 7, as was the case with early assemblies of *P. trichocarpa* (Geraldes *et al*, 2015). An additional 78 SNPs on chr7, sc197, and sc257 exhibited high association with sex but were not below the statistical significance threshold; these did not change the boundaries of the SLR as defined by the 38 SNPs with significant associations (Table S4). The average heterozygosity for males among the 38 SNPs was 0.875 ± 0.218, and the average heterozygosity for females was 0.072 ± 0.148 (mean ± sd), suggesting that the SLR in this species exhibits XY heterogamety (Fig. 2; Tables S5; S6). Amplicons from two loci designed to target areas of this region confirmed these patterns in 32 independent individuals, with an average male heterozygosity of 0.959 ± 0.067 (mean ± sd), and an average female heterozygosity of 0.021 ± 0.088 for 18 SNPs with a MAF > 0.2 (Table S6).

**Figure 1.**
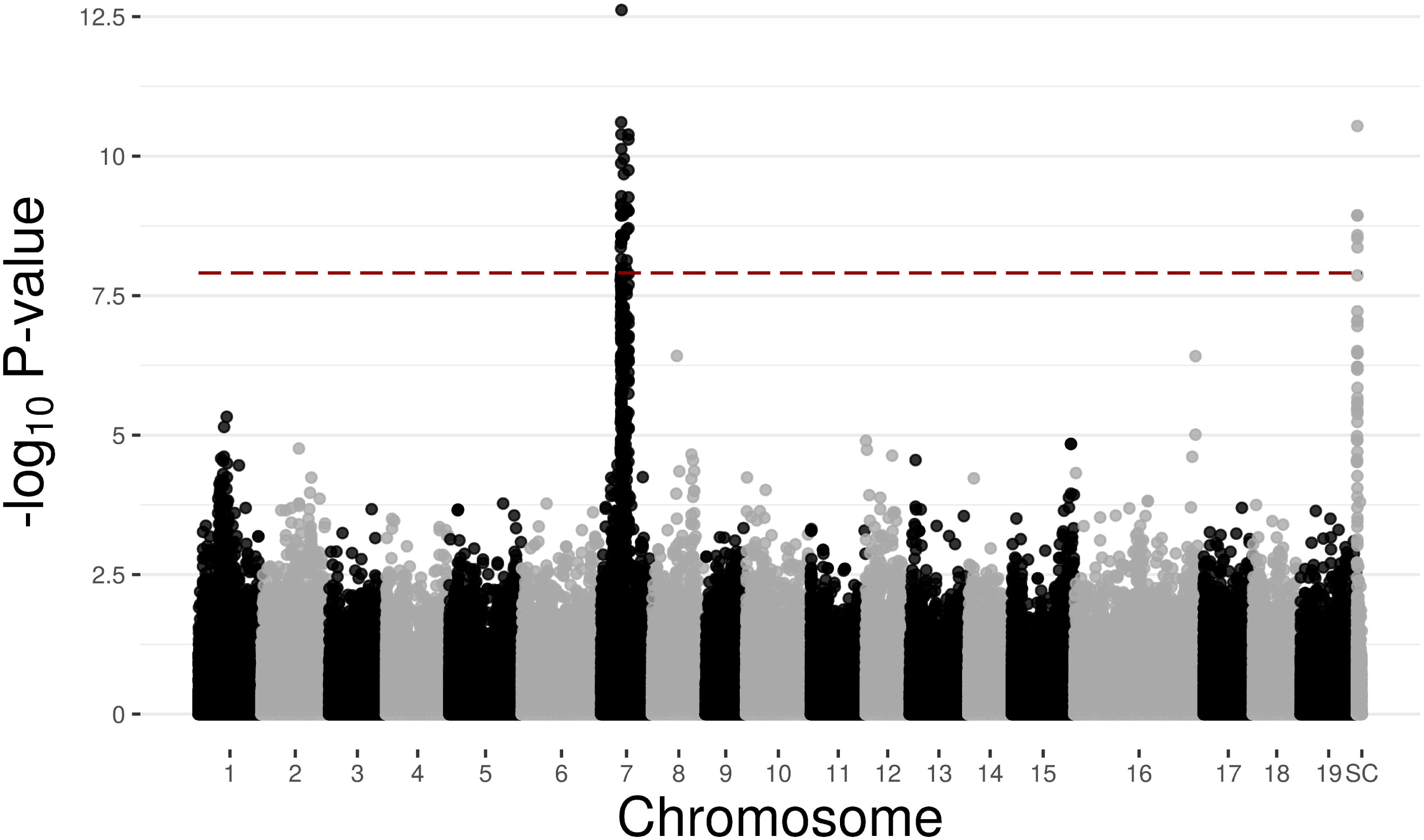
Genome wide associations with sex for 24 male and 24 female *Salix nigra* individuals, expressed in terms of –log_10_ *P*-values. The relative positions of each SNP on each of the 19 chromosomes and unplaced scaffolds (SC) as mapped against the *S. purpurea* genome are shown on the X-axis. Alternating black and grey dots represent different chromosomes. Red dashed line indicates a Bonferroni-adjusted *P*_*FDR*_ = 0.05 (-log_10_ *P*_*FDR*_ = 7.91).

**Figure 2.**
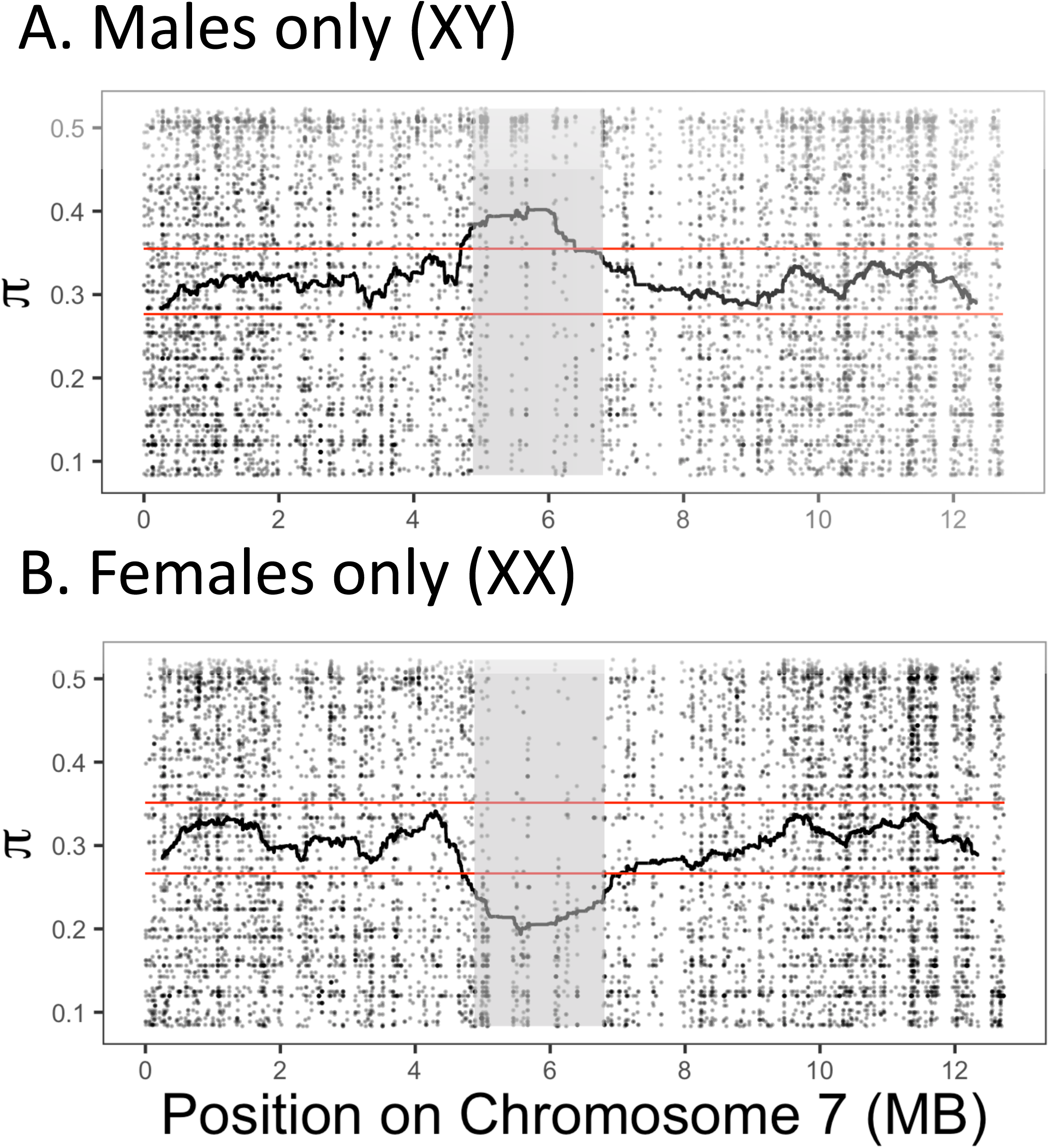
Patterns of nucleotide diversity and heterozygosity on *Salix nigra* chromosome 7. A) Dots represent Nei’s π for all SNPs in males across chromosome 7. Darkness of the dots represent the relative density of overlapping SNPs plotted in the same area on the figure. The black line represents the rolling mean of 5000 base pair windows. Red horizontal lines indicate the 99% and 1% quantiles of non-overlapping 5000 base pair windows across the genome. B) π for all SNPs in females across chromosome 7. Grey area represents the proposed sex-linked region.

Finally, read counts for males in the SLR were slightly lower for males than for females (Figure S1; males: mean=24.8, median=16; females: mean=25.9, median=17; Kruskal-Wallis *χ*^2^=49.2, P<0.001), but they were not half, as would be expected the absence of loci through degeneration of the Y chromosome.

Recombination is suppressed between the X and Y chromosomes in the SLR as shown by the slow decay in linkage disequilibrium exhibited in males (XY; Fig. 3). Females (XX) exhibited faster LD decay than males, but slower than PAR1 and PAR2 (Fig. 3). The distance at which LD decays by half across the SLR in males (1.7 MB) was two orders of magnitude greater than the SLR in females (0.08 MB), PAR1 (males: 0.06MB, females: 0.05MB) and PAR2 (males: 0.03, females: 0.03).

**Figure 3.**
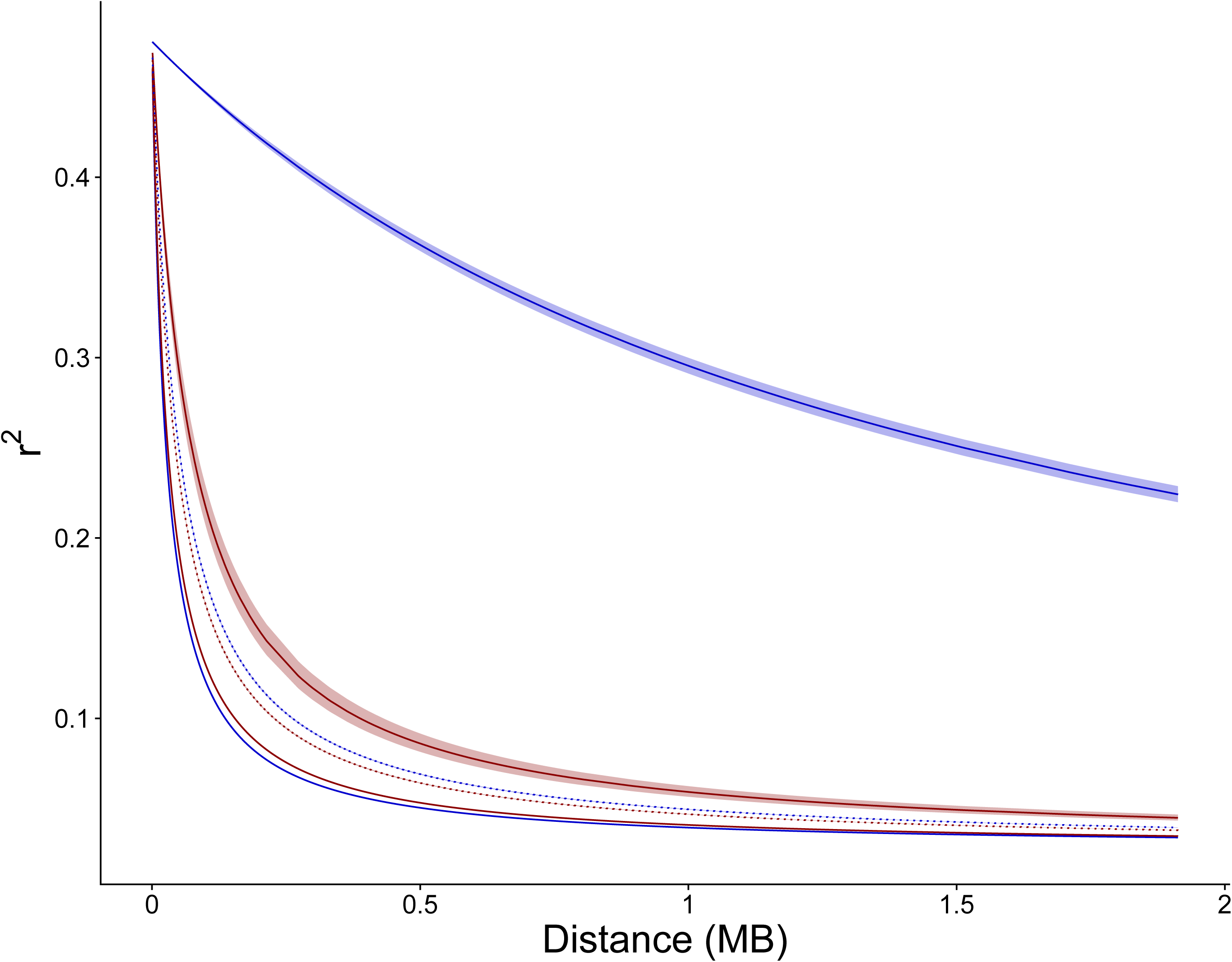
Patterns of linkage disequilibrium decay on *Salix nigra* chromosome 7. LD is expressed as the squared allele frequency correlation (r^2^) between two points, with the distance between points indicated on the X-axis. Blue lines represent LD-decay for chromosome 7 in males and red lines represent LD-decay for chromosome 7 in females. The solid lines represent LD-decay for the SLR, and the recombining regions of chromosome 7 upstream (PAR1) and downstream (PAR2) are represented by dotted and dashed lines, respectively. Ribbons around lines represent the 95% confidence intervals of estimates of r^2^. Note that the 95% CI for PAR1 and PAR2 are plotted, but the intervals around the center line are very small.

Based on homology of our probes to the annotation of the *S. purpurea* genome to which we mapped our reads, our probes targeted 25 genes in the region of chromosome 7 we identified as the SLR plus the two sex-linked scaffolds (Table S7), of which 12 genes carried SNPs significantly associated with sex and an 4 genes carried SNPs with associations greater than any genomic location outside the SLR (Table S7). This set of 16 genes included a homolog to cytokinin oxidase (AT5G56970.1), a possible candidate for presence in the metabolic pathway controlling sex determination (Feng *et al*, 2020). The entire region between 4.88 Mb and 6.81 Mb of chromosome 7 contains 111 genes with *Arabidopsis* orthologs (Table S7) in *S. purpurea*, but the common observation of dramatic restructuring of sex-specific regions when they experience movement in other species (Charlesworth, 2016b; Yang *et al*, 2020; Zhou *et al*, 2020) suggests caution when interpreting too much from homology across this entire interval.

In males the SLR exhibited elevated heterozygosity and pairwise nucleotide diversity compared to the average in autosomes (Fig. 2A), a pattern consistent with XY heterogamety. In females, the SLR exhibited high homozygosity and low pairwise nucleotide diversity compared to the average in autosomes (Fig. 2B), a pattern consistent with lower effective population size of the two X chromosomes carried by females, relative to autosomes. The SLR on the X chromosome also exhibited a large negative deviation in Tajima’s D (mean of 25kb windows = -0.786, 95% CI: -1.270 to -0.232) compared to the autosomes (mean = 0.635, 95% CI: 0.288 to 0.988) and PARs (PAR1 mean = 0.522, 95% CI: 0.156 to 0.852; PAR2 mean = 0.573, 95% CI: 0.196 to 0.938; Fig. 4B & S4), indicating there is an excess of low-frequency polymorphisms in this region. When the population of males was analyzed, the SLR exhibited a large positive deviation in Tajima’s D (Figs. 4B & S4), consistent with the lack of recombination between the X and Y chromosomes.

**Figure 4.**
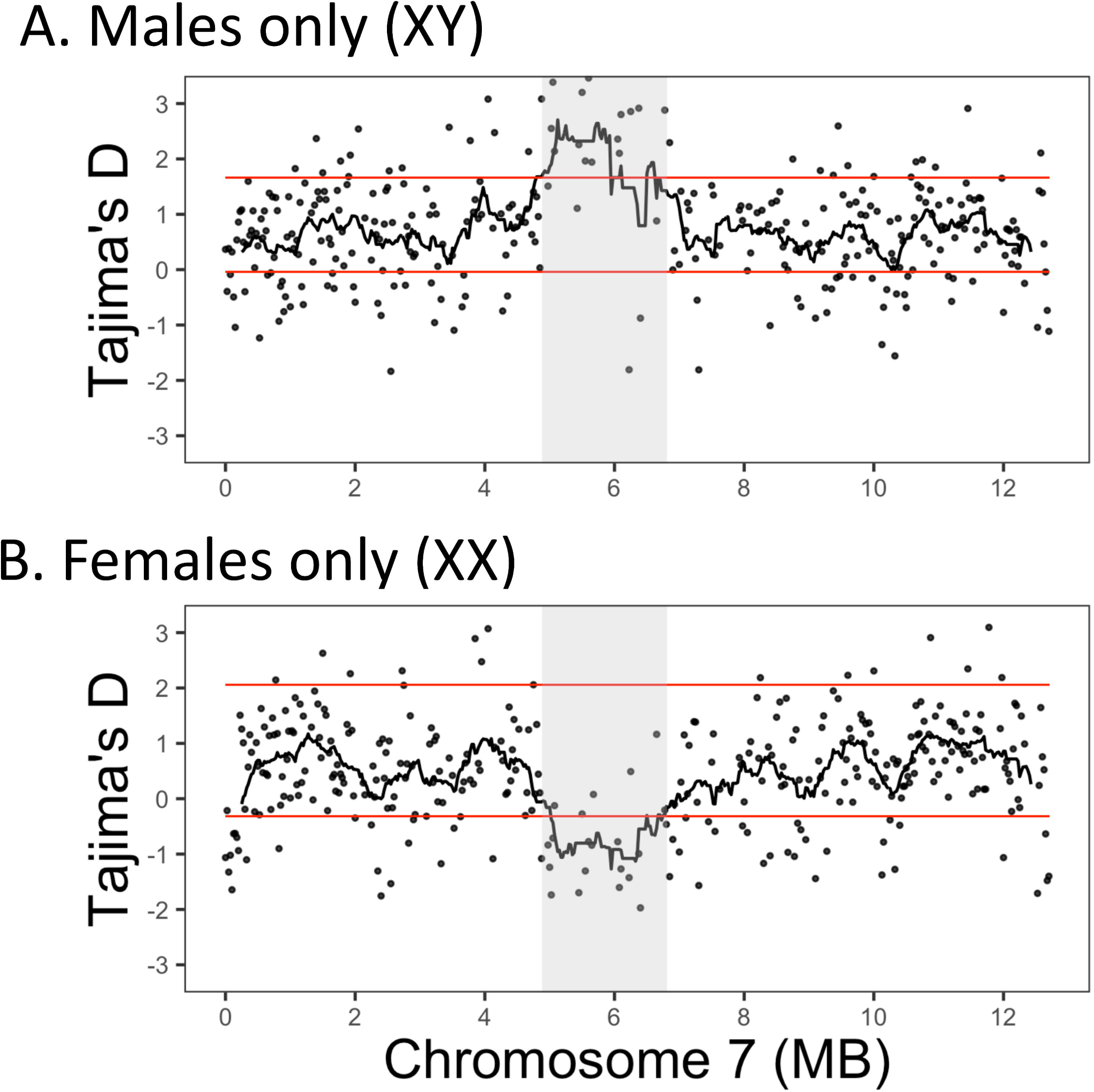
Patterns of Tajima’s D on *Salix nigra* chromosome 7. A) Tajima’s D for 25 kb windows for males (both X and Y chromosomes) across chromosome 7. Dots represent the mean Tajima’s D for non-overlapping 25kb windows across the chromosome. The black line is the rolling mean of 22 25kb windows, the approximate size of the non-recombining sex-linked region (SLR). Red horizontal lines indicate the 99% and 1% quantiles of the Tajima’s D calculated from non-overlapping 25kb windows across the genome including all of chr7. Grey area represents the proposed sex-linked region. B) Tajima’s D for 25 kb windows for females (only X chromosomes) across chromosome 7.

## Discussion

We identified a new sex-linked region (SLR) in willows that exhibits XY heterogamety on chromosome 7 in *Salix nigra*. This result establishes that sex chromosomes in *Salix* have undergone changes in the genomic location as well as patterns of heterogamety. The previously identified sex chromosomes in *Salix* have all been in the subgenus Vetrix, and the location and heterogamety of all of these SLRs was on chromosome 15 ZW (Carlson *et al*, 2017; Hou *et al*, 2015; Pucholt *et al*, 2015; Temmel *et al*, 2007; Zhou *et al*, 2018). The sex chromosomes of poplars (*Populus* spp.), the sister genus to *Salix*, have been mapped to two different SLRs on chromosome 19, with *P. trichocarpa* and *P. tremuloides* exhibiting XY sex determination (Geraldes *et al*, 2015; Pakull *et al*, 2011; Pakull *et al*, 2009) and *P. alba* exhibiting ZW sex determination (Paolucci *et al*, 2010) and a third SLR on chromosome 14 with XY heterogamety in *P. euphratica* (Yang *et al*, 2020). Based on the assumption of strong homology between *S. nigra* and *S. purpurea* on chromosome 7 between 4.88 Mb and 6.81 Mb, the size of the non-recombining SLR in *S. nigra* may be around 2 MB and may contain 111 or more genes. This assumption is not unreasonable given the common chromosome numbers and strong synteny among genomes across *Populus* and *Salix* (Chen *et al*, 2019; Dai *et al*, 2014), however, the sex determination regions in *Populus* and *Salix* are known to be highly dynamic (Yang *et al*, 2020; Zhou *et al*, 2020; Zhou *et al*, 2018). Thus, we expect that these estimates may change dramatically once the SLR is assembled in *S. nigra*. If this estimate is correct, the SLR in *S. nigra* would be intermediate in size between the 6.7 MB SLR in *S. purpurea* (Zhou *et al*, 2020) and the 100kb SLR in *P. trichocarpa* (Geraldes *et al*, 2015).

Dioecy (separate male and female individuals) is found in only 5-6% of angiosperms (Renner and Ricklefs, 1995) and is unevenly distributed across the angiosperm orders (Renner, 2014). Many dioecious species have congeners and sister species that are hermaphroditic, and recent analyses support the hypothesis that dioecy is a key innovation that results in increased diversification within plant clades (Kafer *et al*, 2014). Previous studies have hypothesized that dioecy and the sex determination regions in *Salix* and *Populus* evolved independently (Hou *et al*, 2015), which was formerly supported by the observation that all *Salix* sex chromosomes in the subgenus Vetrix were 15 ZW and most *Populus* sex chromosomes were 19 XY. Recent reports indicate that *Populus* also has sex chromosomes in multiple locations, where a common mechanism of sex determination that includes a cytokinin response regulator (*RR* gene) suggests movement of the sex chromosome within a dioecious lineage (Yang *et al*, 2020). The discovery of the 7 XY SLR in *S. nigra* indicates a previously unknown dynamism in the evolution of sex chromosomes in *Salix*. Our sequence capture library did not include probes for the *RR* gene that is associated with sex determination in both *Populus* (Muller *et al*, 2020) and *Salix* (Zhou *et al*, 2020), so confirmation of the presence or absence of an *RR* homolog or miRNA that targets an *RR* gene in *S. nigra* will require assembly of both the X and Y chromosomes. Such a confirmation of a common sex determination mechanism across both *Salix* and *Populus* would support an alternative hypothesis that dioecy evolved once prior to the split of the genera and the subsequent movement of the sex determination region among dioecious species.

We found that Tajima’s D in the SLR of the X chromosome (chr 7) was negative, whereas Tajima’s D was positive on the autosomes and the PARs (Fig. 4B; Fig. S4B), indicating that minor alleles in the SLR were rare. Several mechanisms may account for different aspects of these patterns including our sampling method, the demographic history of the populations, and selection targeted to genes in the SLR. First, we sampled among individuals from a single small and isolated population in west Texas that exhibited generally high relatedness (Figure S2). Sampling strategy is well known to influence the site frequency spectrum (SFS) and Tajima’s D, and these influences vary depending on the population structure and migration rates among populations (Städler *et al*, 2009). The key implication here is that we should not over-interpret the high value of Tajima’s D in the autosomal sample as a necessarily reflecting a species-wide pattern. Second, theoretical studies indicate that the lower effective population size for the X chromosome than the autosomes (the Ne of the X is 75% of the autosome) results in the X chromosome exhibiting lower values of Tajima’s D than the autosomes during recovery from a population bottleneck (Gattepaille *et al*, 2013). For a bottleneck to produce the strongly divergent patterns between the X chromosome and autosomes (and PARs) observed in *S. nigra*, however, this bottleneck must have been strong, and occurred in the relatively recent past within the very small window of time that is expected to generate both a positive Tajima’s D in autosomes and a negative Tajima’s D on the X chromosome (Gattepaille *et al*, 2013). Interestingly, a similar pattern is apparent when comparing the effects of the recent bottleneck in humans on the SFS for the mitochondrial and autosomal genomes (Fay and Wu, 1999), but the N_e_ of the mitochondrial genome is ¼ that of the autosomes, so differences between Tajima’s D are expected to be much more pronounced and long lived than for differences between the X chromosomes and autosomes. Finally, some form of selection, such as a recent selective sweep or rare recombination associated with purifying selection (Simonsen *et al*, 1995), may have contributed to this pattern in Tajima’s D. These hypotheses can be partially discriminated based on pattern of alleles private to the X and Y chromosomes. A selective sweep on the X without recombination with the Y would maintain and possibly increase the number of alleles unique to either the X or Y, whereas recombination between the X and Y would decrease the numbers of alleles unique to the X or Y. The genotypes of the 38 SNPs significantly associated with sex in the SLR indicated that no alleles were unique to the X or the Y (Table S8), supporting the hypothesis that recombination between the X and Y within the SLR may be contributing to variation on the X. The question remains, however, why are these alleles in low frequency on the X chromosome? If the recombination events have been occurring repeatedly over time, the site frequency spectrum would be expected to reach an equilibrium, with a Tajima’s D that is similar to the remainder of the genome. One possibility is that the low frequency of recombinant alleles is influenced by ongoing background selection on linked sites (Vicoso and Charlesworth, 2006). Greater insight into the processes generating the deviation for a neutral site frequency spectrum on the X chromosome will be possible after assembly of the X & Y chromosomes in *S. nigra*.

We used a targeted sequence capture method to sequence a broad portion of the gene space within the *S. nigra* genome because funds were limited for this project, and this meant that it was only possible to discover variants at loci homologous to our probes. This method allowed us to include 48 individuals within our sample, which increased the power and accuracy to detect alleles associated with sex as well as differences in diversity and Tajima’s D compared to what we would have been able to afford if we had taken a whole genome sequencing approach. Because the density of probes around the SLR was low, we did not capture large regions of the SLR that we may have sampled using whole-genome sequencing (WGS), and this limited our ability to address patterns of diversity across the entire SLR. However, SLRs are notoriously challenging to sequence and assemble, due to the common occurrence of repetitive regions, so a low-coverage WGS approach also may not have recovered a larger portion of the SLR in this novel species. A second limitation of our work was the absence of a reference assembly from *S. nigra*, and instead we relied on a *S. purpurea* reference to map our loci. This is especially limiting when mapping an SLR, which are known to have complex histories of rapid evolutionary change, including translocations, deletions, and inversions (Furman *et al*, 2020).

Nonetheless, all of the SNPs with high association to sex mapped to one of only three locations on the *S. purpurea* reference; either to chromosome 7 or to one of two scaffolds that remain unassembled on the *S. purpurea* v5 assembly. We predict that these scaffolds will map to chromosome 7 as assemblies of *S. purpurea* and *S. nigra* improve. Despite these limitations of the sequence capture approach, we believe these results highlight the utility of this method for inexpensively generating new insights via mapping experiments in novel species.

## Conclusion

This report of a previously unidentified SLR in *S. nigra* on chromosome 7 with XY heterogamety illuminates the dynamic history of the shifting positions and dominance relationships of sex chromosomes in the Salicaceae family, which includes both poplars (*Populus* spp.) and willows (*Salix* spp.). Patterns of Tajima’s D suggest that either *S. nigra* is recovering from a recent population size bottle neck or that the X chromosome was impacted by either a selective sweep or ongoing background selection. It is intriguing to consider that the excess of low frequency alleles may have resulted from selection induced by the recent the movement of the SLRs (Saunders *et al*, 2018) in this plant family, but given that the effects of selection on the SFS are fleeting, this SLR may be sufficiently old that the footprint of the sex chromosome transition on the SFS may no longer be present. This study further supports the Salicaceae as among a small number of taxonomic groups with high dynamism in the turnover of sex chromosomes (Furman *et al*, 2020). Future research should focus on understanding whether additional shifts in the location and dominance relationships of the sex chromosomes have occurred within the family and what mechanisms are driving these changes.

## Acknowledgements

We thank Rao Kottapalli, Pratibha Kottapalli, and Sandra Simon for assistance with library preparation, Julianne Grady for assistance with the amplicon analysis, and Haley Hale for assistance with DNA extraction. We acknowledge Quentin Cronk (UBC, Vancouver) for making available *Salix* genomic resources (supported through funding from Genome Canada, 168BIO). We thank the Dickens County Judge for permission to sample this *S. nigra* population. This research was supported by grants from the US National Science Foundation (1542509, 1542599, 1542479, 1542486) and the National Natural Science Foundation of China (31561123001).

## Author Contributions

B.J.S., S.P.D, and M.S.O designed the experiment; S.P.D., L.B.S., K.K-R., T.Y., J.L., and M.S.O. secured funding for this project. L.B.S., T.M., and J.L. contributed whole-genome data for sequence capture design; B.J.S and S.P.D developed the targeted sequence capture array; M.S.O. collected plant materials; B.J.S., G.F., and N.H. carried out the experiment; B.J.S, S.P.D., G.F., C.H.C., and M.S.O. analyzed the data and prepared figures; B.J.S. and M.S.O. drafted the manuscript, and all authors contributed to revisions of the manuscript.

## Data Accessibility Statement

Upon acceptance of this manuscript the short read data from the targeted sequence capture will be available at the NCBI Sequence Read Archive. All custom scripts used, as well as full Jupyter analysis notebooks are available at https://github.com/BrianSanderson/salix-nigra-slr/, and upon acceptance of this manuscript a DOI will be generated from Zenodo to reference the state of the repository at the time of acceptance.

## Supplemental Information

### Supplemental Figure and Table Legends

**Figure S1.**
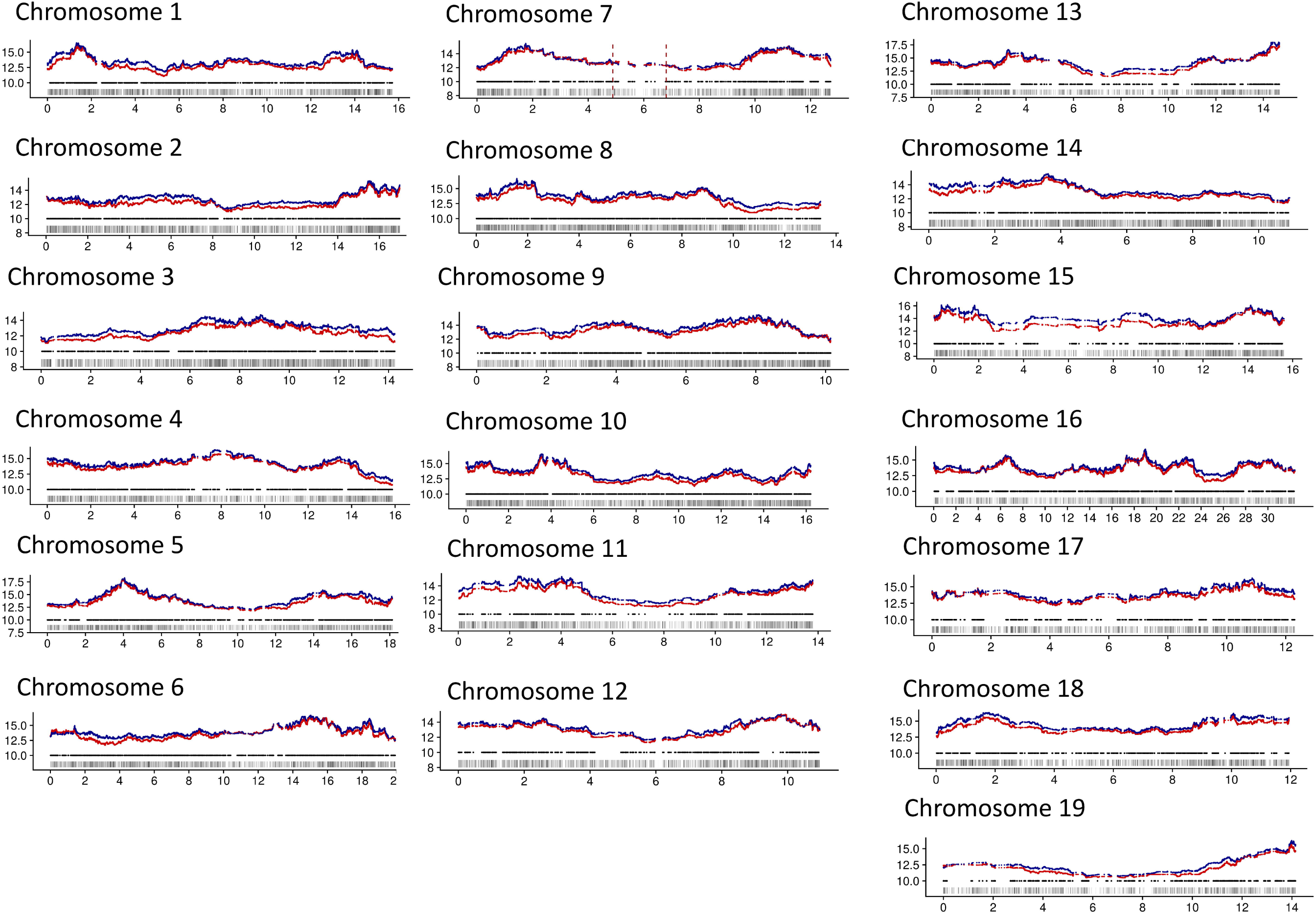
Sequence capture probes and sequencing depth across the 19 chromosomes of Salix nigra. Sequencing reads were mapped to the *Salix purpurea* v5.1 genome. The bottom dashes indicate locations for all genes from the *S. purpurea* v5 annotation. The middle row of black dots are the locations where the sequence capture probes map onto genome using bwa mem. The blue (male) and red (female) dots at the top represent mean read depths for male and female libraries respectively. Depths are plotted as means over 5000 bp windows for all individuals within each sex. Vertical dashed lines on chr7 indicate the proposed location of the SLR. Note that the Y-axis depth values are only meaningful for the blue and red mean depth lines.

**Figure S2.**
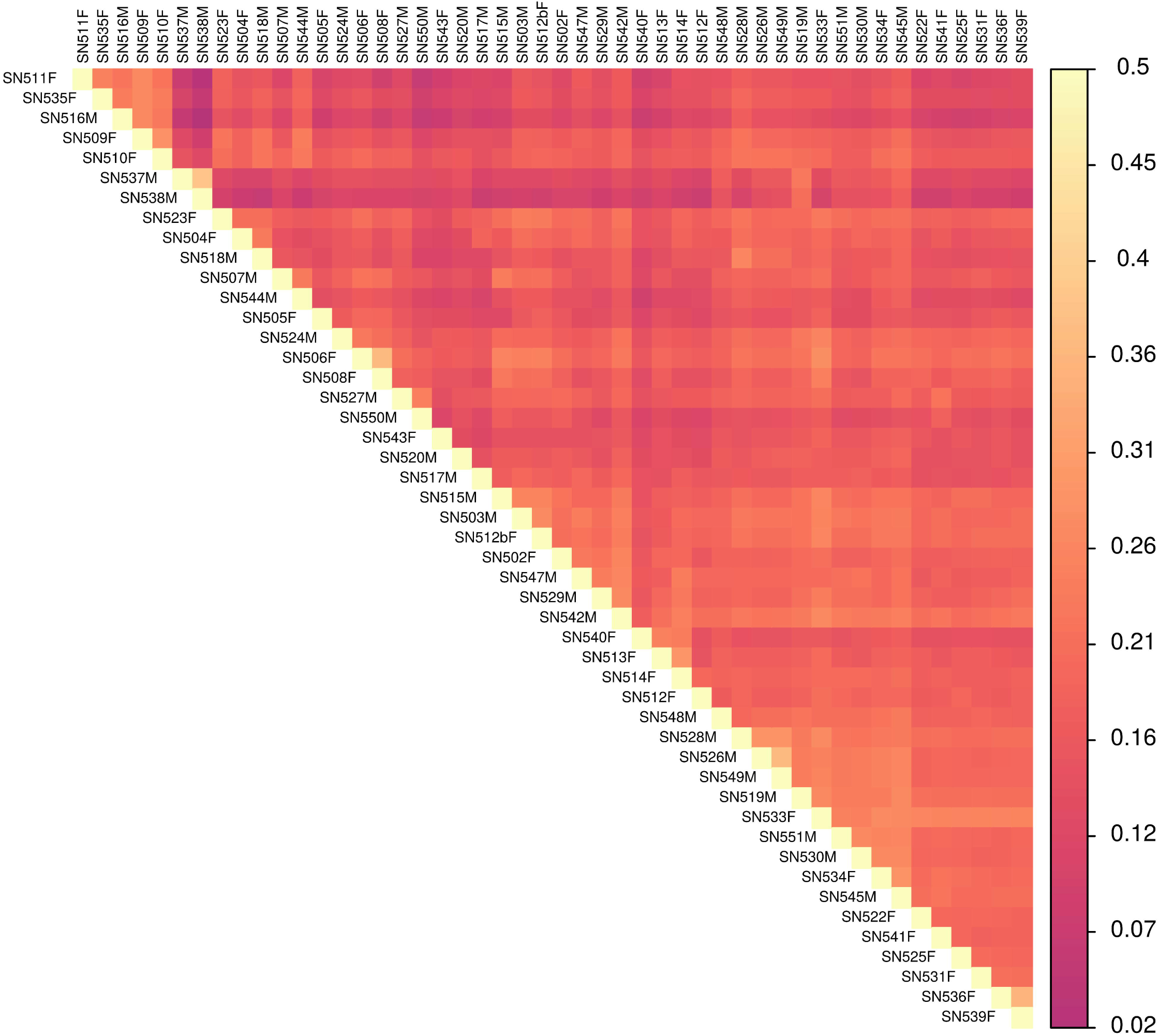
Pairwise kinship estimates between males and female *Salix nigra* in this study. Kinship was estimated using the relatedness2 algorithm in vcftools, which applies the methods used in KING (Manuchaikul et al. 2010).

**Figure S3.**
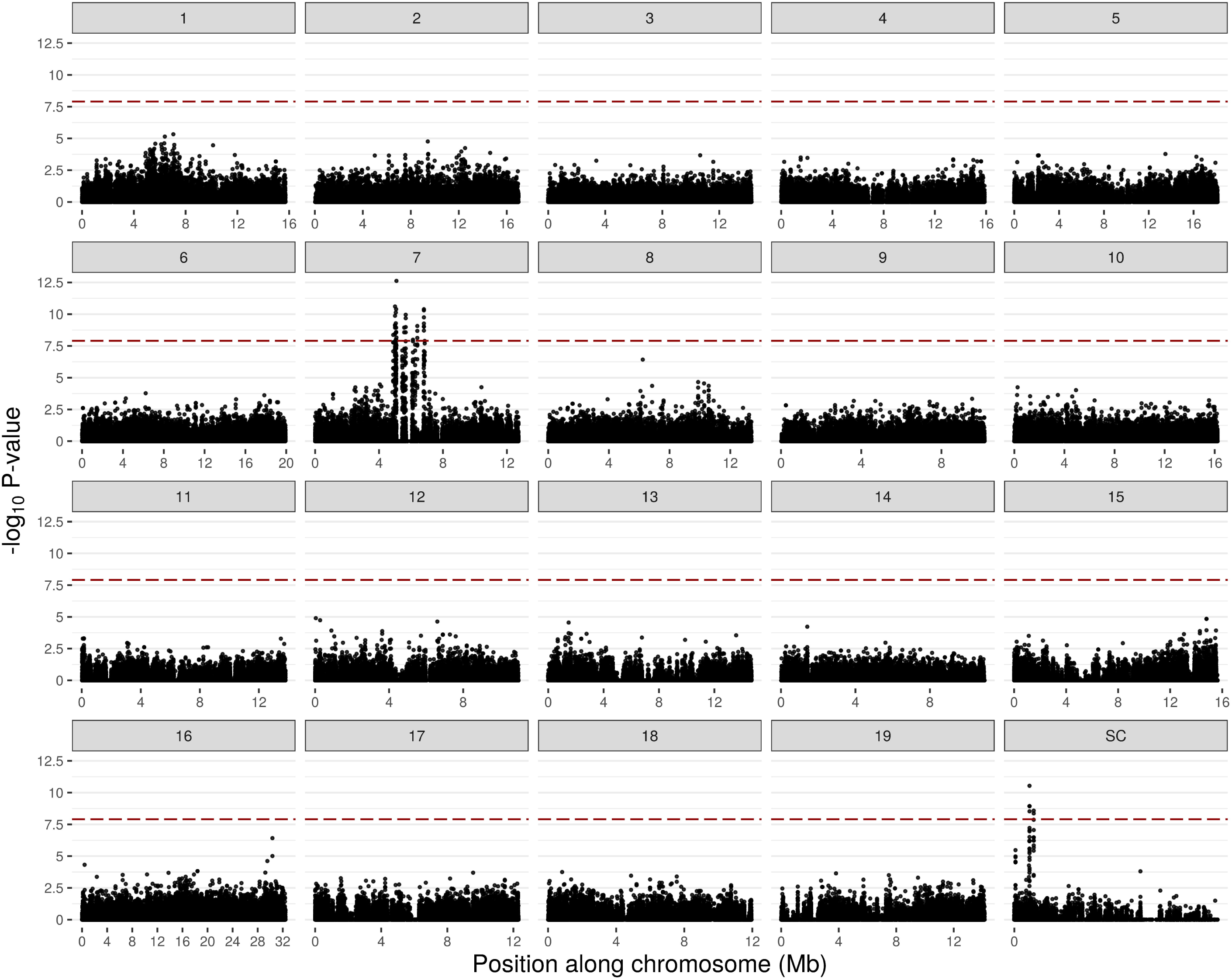
Genome wide associations with sex for 24 male and 24 female *Salix nigra* individuals, expressed in terms of –log_10_ *P*-values plotted for each chromosome. The relative positions of each SNP on each chromosome (numbers in grey titles above each panel) and scaffold (SC) as mapped against the *S. purpurea* genome are shown on the x-axis. The red dashed line indicates a Bonferroni-adjusted *P*_*FDR*_ = 0.05 (-log_10_ *P*_*FDR*_ = 7.91).

**Figure S4.**
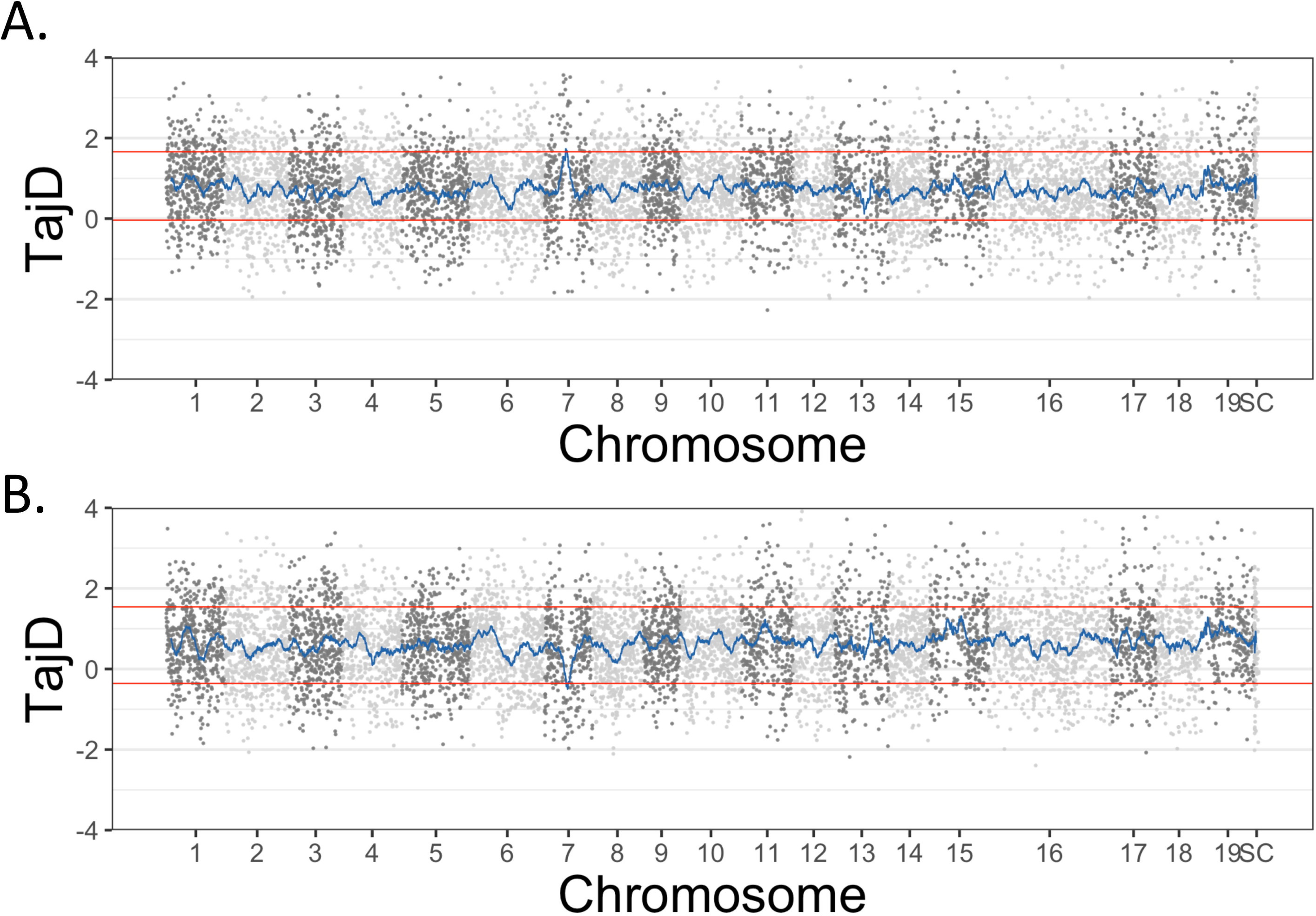
Patterns of Tajima’s D (TajD) across the *Salix nigra* genome. A) Tajima’s D for males only. Dots represent values of Tajima’s D within 25kb across the genome each of the 19 chromosomes and unplaced scaffolds (SC). The blue line is the rolling mean of 22 25kb windows, the approximate size of the non-recombining sex-linked region (SLR). Red horizontal lines indicate the 99% and 1% quantiles of the Tajima’s D calculated from non-overlapping 25kb windows across the genome including all of chr7. B) Tajima’s D for females only.

**Table S1**. Sequence capture probe locations relative to the *S. purpurea* genome V1.1 This array is available from Arbor Biosciences (Ref#170623-30).

**Table S2**. Primers targeting genes in the sex-linked region of *Salix nigra*.

**Table S3**. Read depths for filtered BAM files (on-target) and filtered VCF (whole genome).

**Table S4**. All SNPs on Chr7, Sc197, and sc257 with associations with sex greater than on any other scaffold (highest on other chromosomes was Chr8:6232700, log10P=6.42). SNPs with Bonferroni-adjusted *P*_*FDR*_ > 0.05 (= log_10_ *P*_*FDR*_ > 7.9) are noted in column 2.

**Table S5**. Patterns of heterozygosity for males and females for SNPs with significant sex association on chromosome 7.

**Table S6**. Results of amplicon sequencing of 16 male and 16 female *S. nigra* trees from New York and West Virginia. Results are shown only for loci that had a minor allele frequency > 0.2.

**Table S7**. Genes between 4.88MB and 6.88 MB on chromosome 7 and on scaffold 197 and 257 in the S. purpurea genome based on homology with the *S. purpurea* V5 genome. This region is the location of the non-recombining sex-linked region in *Salix nigra*. **= Genes with at least one SNP with Bonferroni-adjusted *P*_*FDR*_ < 0.05; *= genes with at least one SNP with sex associations greater than on any chromosome except chr7, sc197, and sc257 (but which were not statistically significant). The probes were designed based on the *S. purpurea* V1.1 genome, for which gene names are not always translatable to the V5.1 genome, which was used as a reference for mapping the sequence capture probes. Thus, the presences of probes for each gene in the v5.1 genome was based on the presence of a V5.1 synonym in the Spurpurea_519_v5.1.synonym.txt file prepared by JGI and/or the successful hit after blasting the probe sequences onto the *S. purpurea* V5.1 transcripts.

**Table S8**. Genotypes of all individuals for the 38 significant SNPs on Chromosome 7. A) All genotype calls. 0 = homozygote reference allele, 1 = heterozygote, 2 = homozygote alternate allele, -1 = not called. Because position 6 122 911 had 3 alleles, exact genotypes were called. B) sums across heterozygotes and homozygotes.

## Notes

### Competing Interest Statement

The authors have declared no competing interest.

### Summary of Updates

Many revisions following a round of peer review

